# Minor spliceosome inactivation in the developing mouse cortex causes self-amplifying radial glial cell death and microcephaly

**DOI:** 10.1101/182816

**Authors:** Marybeth Baumgartner, Anouk M. Olthof, Katery C. Hyatt, Christopher Lemoine, Kyle Drake, Nikita Sturrock, Nhut Nguyen, Sahar Al Seesi, Rahul N. Kanadia

## Abstract

Inactivation of the minor spliceosome has been linked to <u>m</u>icrocephalic <u>o</u>steodysplastic <u>p</u>rimordial <u>d</u>warfism type <u>1</u> (MOPD1). To interrogate how minor intron splicing regulates cortical development, we employed *Emx1*-Cre to ablate *Rnu11*, which encodes the minor spliceosome-specific U11 small nuclear RNA (snRNA), in the developing cortex (pallium). *Rnu11* cKO mice were born with microcephaly, caused by death of self-amplifying radial glial cells (RGCs). However, both intermediate progenitor cells (IPCs) and neurons were produced in the U11-null pallium. RNAseq of the pallium revealed elevated <u>mi</u>nor intron retention in the mutant, particularly in genes regulating cell cycle. Moreover, the only downregulated minor intron-containing gene (MIG) was *Spc24*, which regulates kinetochore assembly. These findings were consistent with the observation of fewer RGCs entering cytokinesis prior to RGC loss, underscoring the requirement of minor splicing for cell cycle progression in RGCs. Overall, we provide a potential explanation of how disruption of minor splicing might cause microcephaly in MOPD1.

**Summary Statement:** Here we report the first mammalian model to investigate the role of the minor spliceosome in cortical development and microcephaly.

**List of abbreviations used:** MOPD1=microcephalic osteodysplastic primordial dwarfism type 1; snRNA=small nuclear RNA; cKO=conditional knockout; NPC=neural progenitor cell; RGC=radial glial cell; IPC=intermediate progenitor cell; MIG=minor intron-containing gene

## Introduction

RNA splicing, the removal of non-coding introns from pre-mRNA transcripts, is an essential step in eukaryotic gene expression. This process is performed by one of two distinct spliceosomes, the major or the minor type, both of which consist of five small nuclear RNAs (snRNAs) and associated proteins (Patel and Steitz, 2003). The minor spliceosome is comprised of four unique snRNAs (U11, U12, U4atac, and U6atac) and the U5 snRNA, which it shares with the major spliceosome (Tarn and Steitz, 1996b; Tarn and Steitz, 1996a). In the mouse genome, <0.5% introns, called minor introns, require the minor spliceosome (Levine and Durbin, 2001; Alioto, 2007). In contrast, the major spliceosome, consisting of the U1, U2, U4, U5, and U6 snRNAs, splices the remaining >99.5% of introns, known as major introns (Patel and Steitz, 2003). Minor introns are found embedded in genes that mostly consist of major introns (Alioto, 2007). Thus, proper expression of <u>mi</u>nor intron-containing <u>g</u>enes (MIGs) requires the coordinated action of both the major and minor spliceosomes, thereby making splicing of MIGs relatively inefficient. This inefficiency is further compounded by 100-fold lower expression of the minor spliceosome-specific snRNAs compared to their major snRNA counterparts (Patel et al., 2002). A widely accepted model is that inefficient splicing creates a bottleneck, which acts as an additional layer of regulation of gene expression (Patel et al., 2002). Curation of biological functions of the MIGs does not reveal a common thread that connects the MIGs in regulation of a specific biological process. While the functions executed by the MIGs are disparate (Turunen et al., 2013), the importance of proper minor intron splicing and MIG expression is underscored by the diseases <u>m</u>icrocephalic <u>o</u>steodysplastic <u>p</u>rimordial <u>d</u>warfism type <u>1</u> (MOPD1) and Roifman syndrome, which are characterized by microcephaly and dwarfism (Edery et al., 2011; He et al., 2011; Merico et al., 2015). Both of these conditions are linked to hypomorphic mutation in the U4atac snRNA (Edery et al., 2011; He et al., 2011; Merico et al., 2015). Specifically, in MOPD1 it was shown that the U4atac mutation results in ~90% loss of minor spliceosome activity (He et al. 2011). It is thought that the remaining 10% of minor spliceosome activity allows escape from embryonic lethality, but not the developmental defects observed in the conditions. Thus, despite the constitutive 90% loss of minor spliceosome activity in MOPD1 patients, the cortex and limbs are particularly susceptible to loss of minor spliceosome activity. Here, we explored the role of minor splicing in cortical development. To gain insight into the molecular and developmental processes in corticogenesis that are regulated by minor splicing, we sought to inactivate the minor spliceosome during embryonic development of the mouse cortex. We generated a conditional knockout (cKO) mouse targeting the U11 snRNA, a crucial component of the minor spliceosome that base pairs to the 5’ splice site (5’SS) in the first step in the splicing reaction (Kolossova and Padgett, 1997). Here, we report our findings related to ablation of minor spliceosome activity in the *Emx1*+neural progenitor cells (NPCs) in the pallium that give rise to the majority of the excitatory neurons of the mouse cortex.

## Results

### *Emx1*-Cre-mediated ablation of U11 in the pallium causes microcephaly

The *Rnu11* conditional knockout (cKO) mouse was generated by engineering loxP sites 1090 bp upstream and 1159 bp downstream of the *Rnu11* gene (Fig. 1A, Fig. S1, SI Methods). Successful targeting of the loxP sites was confirmed by long-range nested PCR in targeted embryonic stem cells (Fig. S1B-C) and further validated by the loss of the wild-type (WT) allele in *Rnu11^Flx/Flx^* mice (Figure 1B). *EIIa*-Cre mediated germline recombination generated *Rnu11^WT/KO^* mice, which showed the presence of the KO allele, which was absent in *Rnu11^WT/WT^* genomic DNA (Fig. 1C-E). Quantitative PCR (qPCR) for the WT allele showed 50% reduction in *Rnu11^WT/KO^* mice compared to *Rnu11^WT/WT^* mice (Fig. 1F). Intercrossing *Rnu11^WT/KO^* mice did not yield *Rnu11^KO/KO^* mice (*P*-value (χ^2^)<0.001) (Fig. 1G). Moreover, this cross did not result in any *Rnu11^KO/KO^* embryos as early as embryonic day (E) 9, indicating embryonic lethality occurred prior to this time-point.

**Figure 1.**
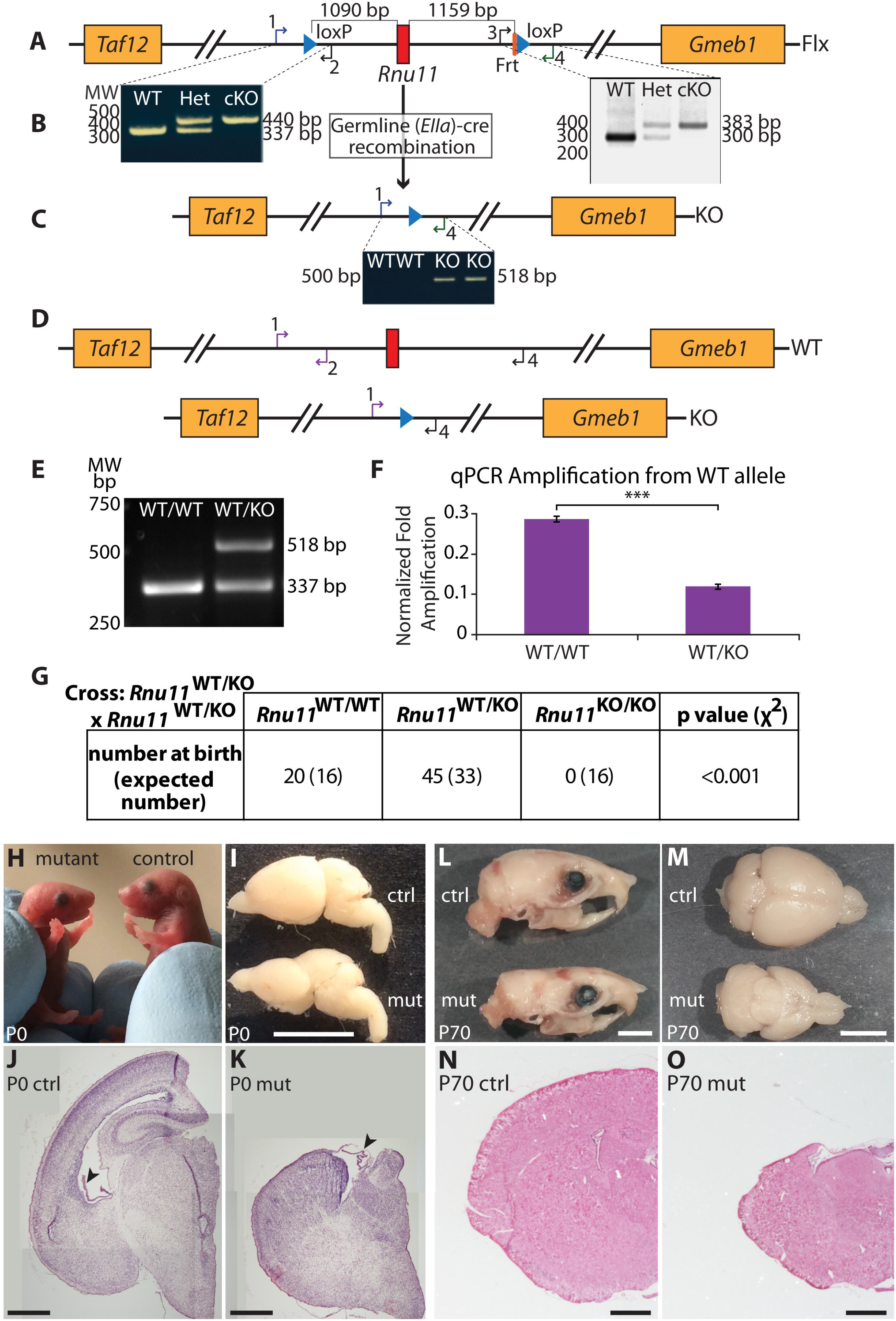
U11 loss in the developing mouse neocortex causes severe microcephaly. (**A**) Schematic of the *Rnu11* floxed (Flx) allele with positions of the loxP sites (blue triangles). (**B**) Agarose gel image showing PCR results detecting the presence of the upstream (left) and downstream (right) loxP sites. (**C**) Schematic of the *Rnu11* knockout (KO) allele that was confirmed by PCR using primers 1 and 4, shown in the gel image (below). (**D**) Schematic showing positions of the primers (1, 2, and 4) on the WT and KO alleles. (**E**) Agarose gel image showing PCR products of the primers shown in (**D**). (**F**) Results of quantitative PCR (qPCR) detecting the WT allele (primers 1 and 2). Significance was determined by a two-tailed student’s t-test; ***=P<0.001. (**G**) Table showing genotype frequency of litters produced from crosses of *Rnu11^WT/KO^* mice, with statistical significance determined by chi-square test. (**H**) Images of P0 *Rnu11^WT/Flx^::Emx1*-Cre^+/-^ (control, right) and *Rnu11^Flx/Flx^::Emx1*-Cre^+/-^ (mutant, left) pups. (**I**) Lateral view of P0 control (ctrl, top) and mutant (mut, bottom) brains. Scale bar=5 mm. (**J-K**) Coronal sections of the P0 brains shown in (**I**) stained with hematoxylin and eosin. Arrowheads indicate the choroid plexis. Scale bar=500 μm. (**L**) Lateral view of skulls from P70 control (top) and mutant (bottom) mice. Scale bar=5 mm. (**M**) Dorsal view of the brains removed from the skulls in (**L**). Scale bar=5 mm. (**N-O**) Coronal sections of the P70 brains in (**M**), stained with hematoxylin and eosin. Scale bar=1 mm.

To circumvent embryonic lethality, we employed *Emx1*-Cre mice to specifically ablate *Rnu11* in the NPCs of the developing cortex (pallium) (Gorski et al., 2002). Ablation of U11 in NPCs in the pallium resulted in profound microcephaly at birth in *Rnu11^Flx/Flx^:.Emx1*-Cre^*+/-*^ mutant mice (Fig. 1H-I). Hematoxylin and eosin (H&E) staining of coronal sections of the P0 mutant brain showed collapse of the cortex and absence of the hippocampal structures (Fig. 1J-K). Approaching the age of weaning (P21), mutant mice showed slower weight gain (Fig. S2A), indicating that they struggled to transition to solid food. Notably, previous reports have shown weak LacZ Cre reporter expression in the first branchial arch in *Emx1*-Cre E10.5 embryos; this structure gives rise to the developing jaw (Gorski et al., 2002; Inman et al., 2013). Indeed, we observed expression of a tdTomato Cre reporter in the jaw of P0 control and mutant mice (Fig. S2B-C); this expression persisted in both the masseter and temporalis muscles of P21 control and mutant mice (Fig. S2D-E’) (Madisen et al., 2010). Anticipating that the mutant mice would have difficulty masticating solid food, we wet their food daily starting at P19, which allowed most mutant mice to survive to adulthood, with some surviving up to 1.5 years (Fig. S2F). The cortex of the adult (P70) mutant mice appeared to have undergone further reduction in size, relative to that seen at P0 (Fig. 1L-O). Interestingly, the adult mutant mice could breed successfully, although the females did not take care of their pups. Despite providing wet food for the mutant mice, *Rnu11^Flx/Flx^::Emx1*-Cre^*+/+*^ mice still died within a week of weaning (Fig. S2G, dotted line). Our findings suggest that homozygosity of *Emx1*-Cre may result in increased recombination in the first branchial arch, failure to transition to solid food, malnourishment, and early death, even when wet food is provided.

### Loss of U11 snRNA results in cell death

Next, we investigated when *Emx1*-Cre-mediated ablation of *Rnu11* resulted in loss of U11 snRNA expression in the *Emx1*-expressing domain of the pallium. Since *Emx1*-Cre is first expressed at E9.5 (Cocas et al., 2009), we performed section *in situ* hybridization (ISH) for U11 snRNA on E10, E11, E12, E13, and E14 pallium sections. Compared to the U11 expression (purple signal) in the E10 control (*Rnu11^WT/Flx^::Emx1*-Cre^*+/-*^) pallium (Fig. 2A), we observed a reduction in U11 signal in the E10 mutant (*Rnu11^Flx/Flx^::Emx1*-Cre^*+/-*^) pallium (white patches, Fig. 2B&B’). The area of this U11-null domain was larger in the E11 mutant pallium (Fig. 2D&D’), and by E12, the majority of cells in the *Emx1*-Cre domain lacked U11 expression (Fig. 2F&F’). We observed scattered cells that retained U11 expression at this point (Fig. 2F’), which may represent either inefficient *Emx1*-Cre recombination at this time-point or migrating interneurons produced in the ventral telencephalon, where *Emx1*-Cre is not active (Gorski et al., 2002; Bartolini et al., 2013). Overall, there was no observable morphological difference between mutant and littermate control palliums at E10, E11, and E12 (Fig. 2A-F). At E13, we observed persistence of the U11-null domain along with collapse of the lateral ventricle in the mutant telencephalon (Fig. 2H&H’). At E14, the thickness of the mutant pallium was drastically reduced (Fig. 2I-J’). We also observed the presence of both U11-null and scattered U11+ cells in the tissue at this time-point (Fig. 2J&J’).

**Figure 2.**
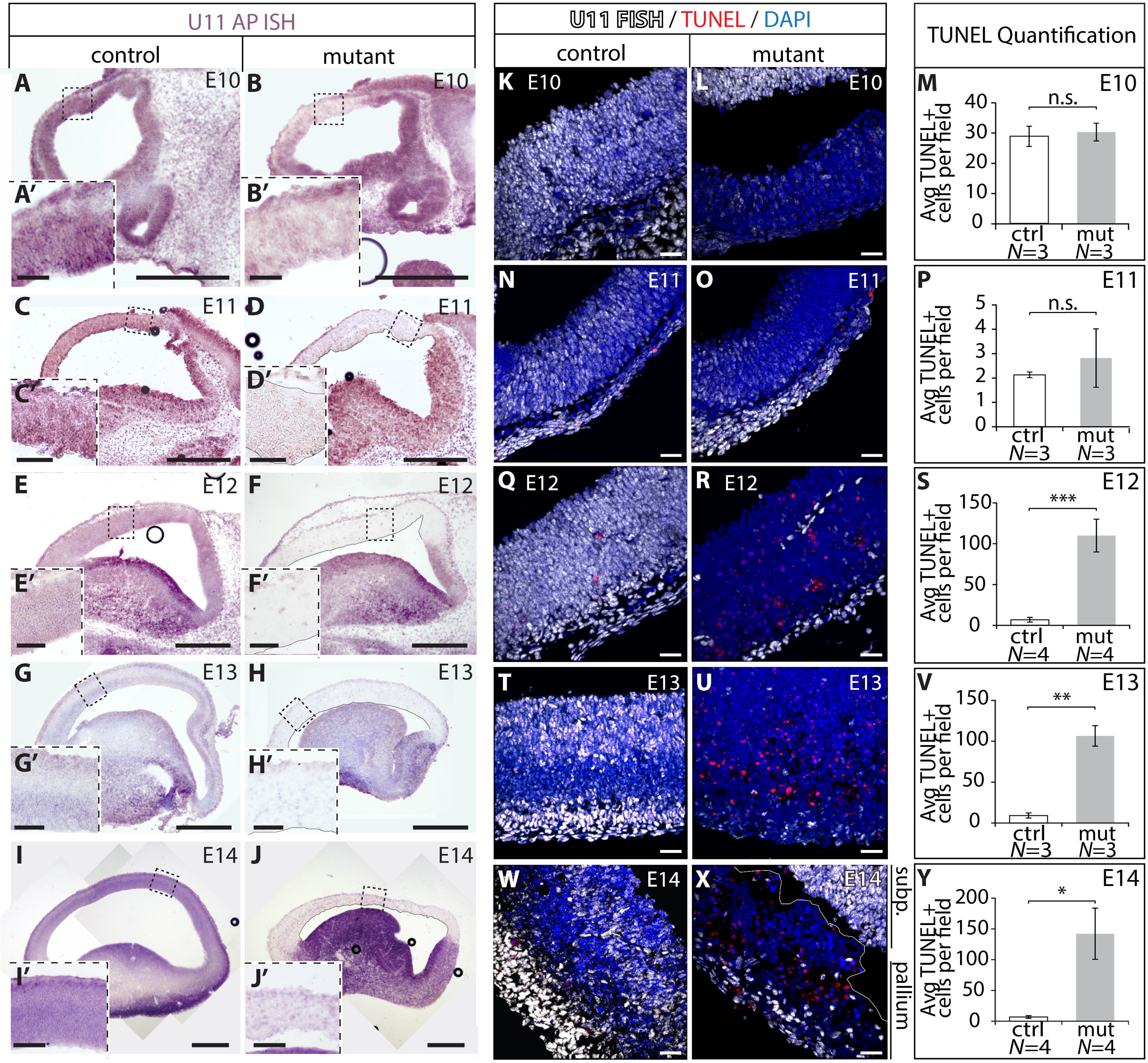
U11 loss does not result in immediate cell death. (**A-J**) U11 expression (purple signal) by *in situ* hybridization (ISH) in sagittal sections of the telencephalon from control (left panels) and mutant (right panels) E10, E11, E12, E13, and E14 embryos. Scale bars=500 μm. Insets (**A’-J’**) show higher magnification of the boxed regions. Scale bars=200 μm. (**K-Y**) Fluorescent ISH (FISH) signal for U11 (white) combined with TUNEL (red) on control (left panels) and mutant (middle panels) pallium sections from E10, E11, E12, E13, and E14 embryos, with bar graphs showing quantification of TUNEL+ cells (right panels). Nuclei are marked by DAPI (blue). Avg=average. Scale bars=30μm. Statistical significance was determined by two-tailed student’s t-tests. N.s.=not significant; *=*P*<0.05; **=*P*<0.01; ***=*P*<0.001.

Reduction in the size of the mutant pallium by E14 suggested either a loss of cells or impaired cell production. To address this issue, we first investigated whether cell death was contributing to the loss of cells in and reduction of the size of the pallium. We performed fluorescent ISH (FISH) to detect loss of U11 followed by terminal dUTP nick-end labeling (TUNEL) to detect apoptotic cells in E10 – E14 pallium sections. Results showed sporadic loss of U11 snRNA in the E10 and E11 mutant sections without a significant increase in TUNEL+ cells (Fig. 2K-P, Fig. S3A-D’’). Onset of cell death in the U11-null pallium was observed at E12, with a statistically significant increase in TUNEL+ cells in the mutant compared to littermate controls (Fig. 2Q-S, Fig. S3E-F”). This pattern of cell death was further exaggerated at E13 followed by reduction at E14 (Fig. 2T-Y, Fig. S3G-J”).

### Loss of U11 snRNA results in reduction of NPCs but not neurons

The cell type being lost in the mutant was identified by immunofluorescence (IF) with antibodies for Ki67 and NeuN to identify proliferating NPCs and neurons, respectively (Wolf et al., 1996; Scholzen and Gerdes, 2000). While there was no difference in the percentage of Ki67+ and NeuN+ cells at E12 (Fig. 3A-B, 3G), at E13 there was a precipitous decline in the percentage of Ki67+ NPCs in the mutant pallium compared to controls (Fig. 3C-D, 3G). This shift could be due to either loss of NPCs or an increase in the number of NeuN+ neurons in the mutant at this time-point. However, there was no change in the absolute number of neurons between the mutant and control at E13 (Fig. S4A), indicating that the shift in Ki67+ NPCs is caused by NPC loss in the E13 mutant pallium. As expected, based on depletion of the NPC population by this time-point, there were significantly fewer neurons in the E14 mutant pallium compared to the control (Fig. 3E-G, Fig. S4A). However, the average number of NeuN+ cells appeared to plateau between E13 and E14 in the mutant pallium (Fig. S4A). One-way ANOVA revealed significant variation in the number of neurons across these three time-points (*P*=0.012). Post hoc Tukey multiple comparison test identified a significant increase in neurons between E12 and E13, but no significant change in the number of neurons between E13 and E14 (Fig. S4B). Together, these data suggested that the cycling U11-null NPCs were most likely dying, while both U11-null NPCs that produce neurons and U11-null neurons managed to escape this fate. Furthermore, U11 FISH in conjunction with IF for Ki67 and NeuN confirmed the presence of both U11-null NPCs and U11-null neurons (Fig. S5). However, the population of U11-null NPCs, and not U11-null neurons, underwent progressive reduction in the mutant pallium.

**Figure 3.**
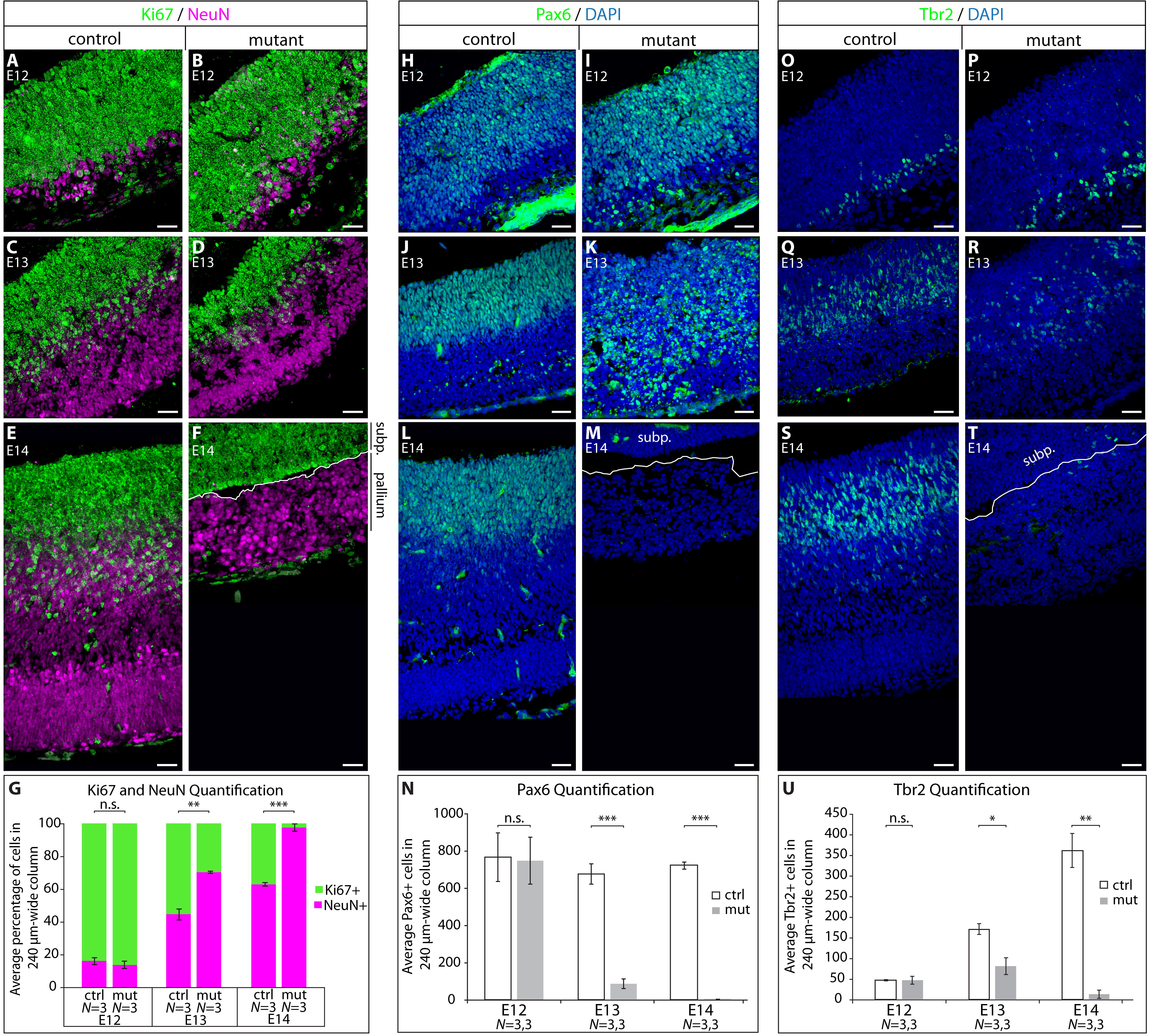
U11 loss results in depletion of NPCs, but not neurons. (**A-F**) Immunofluorescence (IF) for Ki67 (green) and NeuN (magenta) on sagittal sections of the control (left panels) and mutant (right panels) pallium across E12, E13, and E14. (**G**) Bar graph showing quantification of %Ki67+ cells (green) and %NeuN+ cells (magenta), of all labeled cells, within a given 240 μm-wide column of the E12, E13, and E14 pallium. (**H-M**) IF for Pax6 (green), with nuclei stained by DAPI (blue), on sagittal sections of the control (left panels) and mutant (right panels) pallium across E12, E13, and E14. (**N**) Bar graphs showing quantification of the average number of cells with Pax6+ nuclei within a given 240 μm-wide column of E12, E13, and E14 pallium. (**O-T**) IF for Tbr2 (green) on sagittal sections of the control (left panels) and mutant (right panels) pallium across E12, E13, and E14, with nuclei marked by DAPI (blue). (**U**) Bar graph showing quantification of the average number of cells with Tbr2+ nuclei within a given 240 μm-wide column of E12, E13, and E14 pallium. Statistical significance for quantifications was determined by two-tailed student’s t-tests. Subp.=subpallium; n.s.=not significant; *=*P*<0.05; **=*P*<0.01; ***=*P*<0.001. Scale bars=30 μm.

### Self-amplifying RGCs require the minor spliceosome

The precipitous loss of Ki67+ NPCs led us to investigate the type of NPCs being lost in the U11-null pallium. In corticogenesis, there are two types of NPCs: radial glia cells (RGCs) and intermediate progenitor cells (IPCs) (Englund et al., 2005). We interrogated the number of RGCs by IF for the nuclear marker Pax6 (Gotz et al., 1998). This revealed no change in the number of nuclear Pax6+ RGCs at E12 (Fig. 3H-I, 3N). At E13, there was a significant reduction in the number of RGCs with nuclear Pax6 signal in the mutant compared to the control (Fig. 3J-K, 3N). However, widespread cytoplasmic Pax6 expression, which was rarely observed in control sections, was observed in the mutant (Fig. 3J-K). This cytoplasmic staining was primarily observed around condensed nuclei, suggesting that the cytoplasmic Pax6 expression was occurring in dying cells. This cytoplasmic Pax6 expression localized with cleaved caspase 3 (CC3), a marker for cell death (Faleiro et al., 1997) (Fig. S6D-D’’’). Therefore, only cells with nuclear Pax6 expression were quantified. At E14, there were few Pax6+ RGCs in the mutant, which is consistent with the fact that only 5% of the cells were Ki67+ at this time-point (Fig. 3G, 3L-N).

In the U11-null pallium, loss of NPCs occurred without the corresponding reduction in neurons at E13 (Fig. 3G, Fig. S4A). Of the two NPC types in the pallium, IPCs are the source of the majority of neurons in cortical development (Kowalczyk et al., 2009). Therefore, we performed IF using the IPC-specific marker Tbr2 (Englund et al., 2005), which revealed no significant difference in the number of IPCs in the E12 mutant compared to the control (Fig. 3O-P, 3U). At E13, we observed a significant reduction in the IPC population in the mutant relative to the control (Fig. 3Q-R, 3U). By E14, few IPCs were identified in the mutant, which is in agreement with the observed trend of Ki67+ NPC loss at this time point (Fig. 3G, 3S-U).

The trends of RGC and IPC loss during mutant pallium development were not similar: While there was a severe decline in the number of RGCs at E13 compared to E12, there was a modest increase in the number of IPCs at this time-point (Fig. 3N&U). One-way ANOVA revealed significant variation among these three time-points in the mutant (Pax6: *P*=0.001; Tbr2: *P*=0.046). Specifically, the difference between the number of RGCs at E12 and E13 was significant by Tukey test (Fig. S4C). However, for IPCs, there was no significant change between E12 and E13; a significant decline in IPC number was only observed between E13 and E14 (Fig. S4D). The observation that the decline in RGCs occurred earlier within the mutant pallium than the decline in IPCs suggested that the majority of cells dying between E12 and E13 were RGCs.

### RNAseq revealed that inactivation of the minor spliceosome does not affect expression of most MIGs

To understand the underlying molecular defects that led to the loss of self-amplifying RGCs, we performed RNAseq on control (*N*=5) and mutant (*N*=5) palliums from E12 embryos. At this time-point U11 loss occurs throughout the entire *Emx1*-Cre domain without major histological changes, such as shifts in NPC number (Figs 2F, 3G). Confirmation of the dissection of the pallium was reflected by the expression of *Emx1* in the control (19.1 <u>f</u>ragments <u>p</u>er <u>k</u>ilobase per <u>m</u>illion mapped reads [FPKM]) and mutant (20.3 FPKM). Expression of *Dbx1*, which marks the pallial-subpallial boundary (Teissier et al., 2010), and *Lhx6*, which is expressed in the medial ganglionic eminence (MGE) (Grant et al., 2012), were expressed below 1 FPKM in both the control and mutant samples (Supplementary Data File 1). Together, these results confirm the integrity of our dissection. *Rnu11* expression was reduced by 2.95 fold in the mutant compared to the control, which was further confirmed by quantitative reverse transcriptase-PCR (qRT-PCR) (Fig. 4B, Supplementary Data File 1). The incomplete loss of U11 expression in the mutant likely reflects (1) contamination of non-*Emx1*-lineage cells of the meninges and blood vessels (Fig. 2F&F’, Fig. 4A inset) (Siegenthaler and Pleasure, 2011); (2) contamination from non-*Emx1*-lineage cells migrating into the pallium (Fig. 2F&F’) (Bartolini et al., 2013); and/or (3) incomplete recombination by *Emx1*-Cre. Regardless, the mutant samples showed a clear reduction in U11 expression compared to the control, which was in agreement with the ISH analysis (Fig. 2E-F’). Next, we interrogated the expression of all protein-coding genes with ≥1 FPKM in at least one condition. Overall, there was an increase in the number of genes upregulated (74; ≥2-fold change [FC], *P*≤0.01) relative to those downregulated (11; ≥2-FC, *P*≤.01) in the mutant (Fig. 4A, SI Methods, Supplementary Data File 1). DAVID analysis (Huang da et al., 2009a; Huang da et al., 2009b) performed on the upregulated protein-coding genes revealed significant enrichment by Benjamini score (*P*<0.05) of “intrinsic apoptotic signaling pathway in response to DNA damage by p53 class mediator” (Table S1), which is consistent with the cell death observed at this time-point (Fig. 2R-S). Since ablation of *Rnu11* is expected to inactivate the minor spliceosome, we next investigated the expression of its targets, i.e., the MIGs. Surprisingly, the vast majority of MIGs were not differentially expressed (Fig. 4A). Only one MIG, *Spc24*, showed >2-fold downregulation in the mutant (Fig. 4A, arrow), which was independently confirmed by qRT-PCR (Fig. 4C).

**Figure 4.**
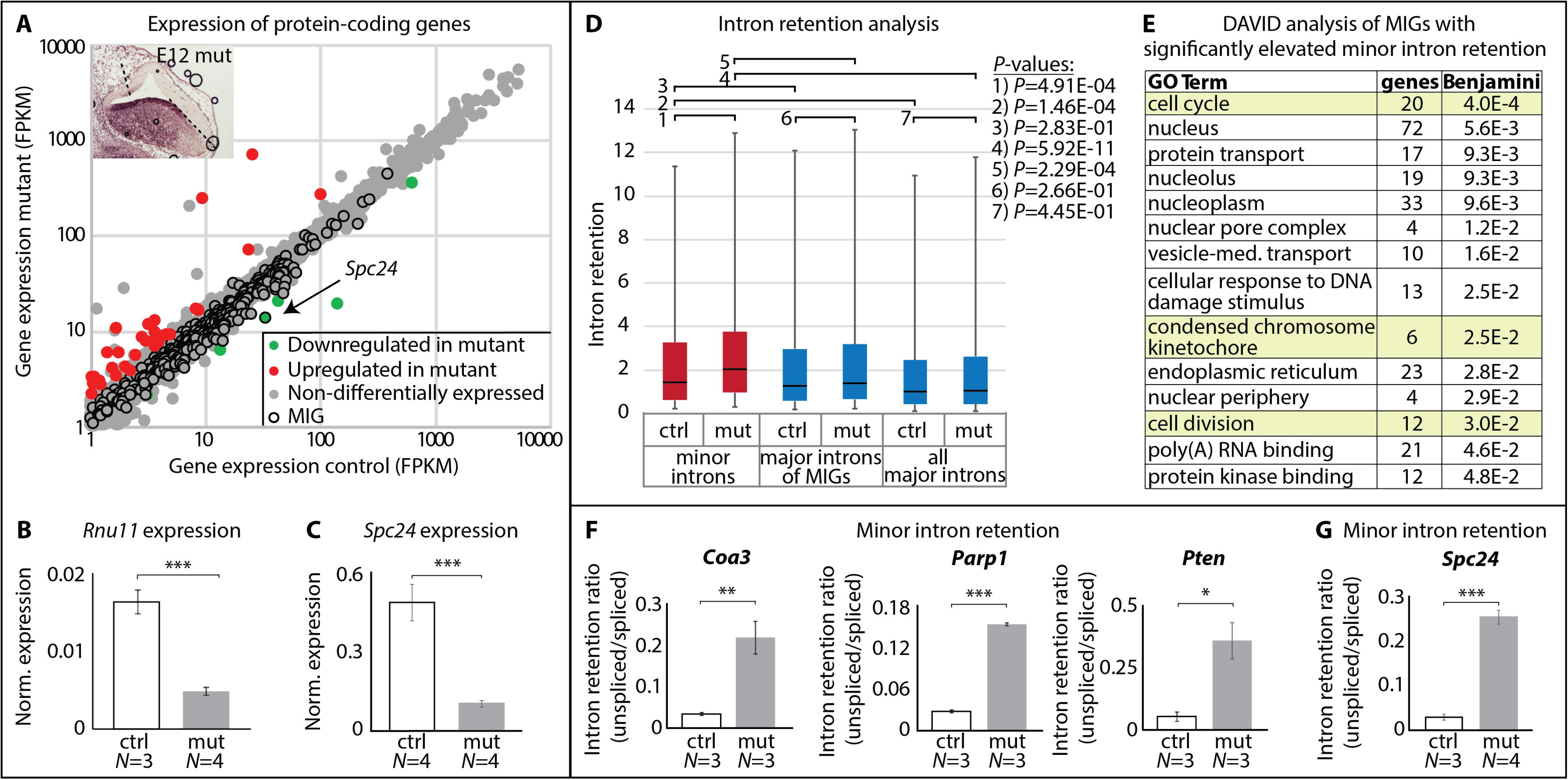
U11 loss in the mutant pallium causes elevated minor intron retention. (**A**) Scatterplot depicting the expression of protein-coding genes ≥1 FPKM in the control (x-axis) and mutant (y-axis) E12 pallium. Inset shows the U11-null domain by ISH on sagittal sections in the E12 mutant pallium, where the dashed lines mark the dissected region. (**B-C**) Normalized expression, determined by quantitative reverse-transcriptase PCR (qRT-PCR) on E12 control and mutant pallium, for *Rnu11* (**B**) and *Spc24* (**C**) (SI Methods). (**D**) Boxplots indicating 5^th^-95^th^ percentile of intron retention for minor introns (red) and major introns (blue) in the control and mutant pallium. The horizontal black lines indicate the median for each condition. Statistical significance was calculated by Wilcoxon rank sum tests. (**E**) Results of DAVID analysis of the MIGs with statistically significant elevated minor intron retention, showing GO Terms and Benjamini values. Functions related to cell cycle are marked in yellow. (**F-G**) Intron retention ratios for the minor introns of the MIGs *Coa3, Parp1*, and *Pten* (**F**) and *Spc24* (**G**) in E12 control and mutant pallium, as determined by qRT-PCR (SI Methods). Norm.=normalized. Statistical significance for qRT-PCR (**B-C, F-G**) was determined by a two-tailed student’s t-tests. *=*P*<0.05, **=*P*<0.01, ***=*P*<0.001. See also Table S2 and Supplementary Data Files 1&2.

### Inactivation of the minor spliceosome affects minor intron splicing

We next analyzed the RNAseq data to investigate minor intron splicing, which we measured via minor intron retention (Fig. S7, SI Methods). Consistent with the inactivation of the minor spliceosome, minor intron retention was significantly higher in the U11-null tissue than the control (Fig. 4D1). We also found that, in the control, minor introns were significantly retained compared to all major introns (Fig. 4D2), which is in agreement with the current model of inefficient minor intron splicing (Patel et al., 2002). However, in the control, comparison of minor intron retention to the retention of major introns found in MIGs did not reveal a significant difference (Fig. 4D3). This suggests that the splicing of major introns within MIGs is affected by the rate of minor intron splicing. In the mutant, minor introns were significantly retained compared to the major introns found in MIGs (Fig. 4D5). Moreover, in the mutant, minor introns were significantly retained compared to all major introns in the mouse genome (Fig. 4D4). Finally, we did not observe a significant difference in major intron retention between the control and mutant (Fig. 4D6, 4D7).

While minor intron retention was significantly elevated in the mutant compared to the control, there was a wide range in the level of retention across the minor introns. Comparison of the minor intron retention values in individual control and mutant samples revealed that 172 of the 408 interrogated minor introns showed significantly elevated retention (*P*≤0.05) in the mutant (Supplementary Data File 2). The elevated minor intron retention for three MIGs (*Coa3, Parp1*, and *Pten*) was confirmed by qRT-PCR (Fig. 4F, SI Methods). Since many minor introns are found within the open reading frame (ORF) of MIGs (3), retention could result in frameshifts, the introduction of premature stop codons, and/or the production of aberrant protein, ultimately disrupting MIG function. To identify which biological processes would be affected in the mutant, we submitted the 169 MIGs with elevated minor intron retention to DAVID (Huang da et al., 2009b; Huang da et al., 2009a). Functions enriched by these MIGs included cell cycle, condensed chromosome kinetochore, response to DNA damage stimulus, and cell division (Fig. 4E, Table S2). Moreover, *Spc24*, the only MIG with significant >2-fold downregulation in the mutant pallium, showed significantly elevated minor intron retention, which was further confirmed by qRT-PCR (Fig. 4A&G; Table S2; Supplementary Data Files 1&2). In all, disruption of minor intron splicing most likely affected cell cycle.

### U11-null NPCs experience cell cycle defects

Spc24 is required to establish and maintain kinetochore-microtubule attachment during mitosis (McCleland et al., 2004). Therefore, loss of *Spc24* function, and/or of the other MIGs involved in cell cycle, would likely result in delayed or failed mitosis and cytokinesis. Since there was no difference in the number of RGCs or IPCs at E12 in the U11-null pallium compared to the control (Fig. 3H-I, 3N, 3O-P, 3U), we sought to determine if there were any observable changes in NPCs at the different stages in cell cycle. For this, we performed IF with antibodies against Aurora B, which marks mitotic cells, and Citk, which marks the cytokinetic midbody (Vader et al., 2006; Paramasivam et al., 2007). Our results showed no difference in the number of Aurora B+ cells in the mutant compared to the control at E11 or E12 (Fig. 5A-D, 5I). In contrast, there was a significant reduction of Aurora B+ cells in the mutant at E13 and E14 compared to the control (Fig. 5E-I), which is consistent with the Ki67 staining (Fig. 3A-G). In the case of Citk, no significant change was observed in the mutant at E11 (Fig. 5J-K, 5R). However, significant reduction in the number of Citk+ cells was observed in the mutant at E12 compared to control (Fig. 5L-M, 5R), which continued at E13 and E14 (Fig. 5N-R). This finding suggested that fewer U11-null NPCs were entering cytokinesis, which is in agreement with the downregulation and increased minor intron retention of *Spc24*, along with the elevated minor intron retention of 19 other cell cycle-regulating MIGs (Fig. 4A, 4C, 4E, 4G; Table S2; Supplementary Data Files 1&2).

**Figure 5.**
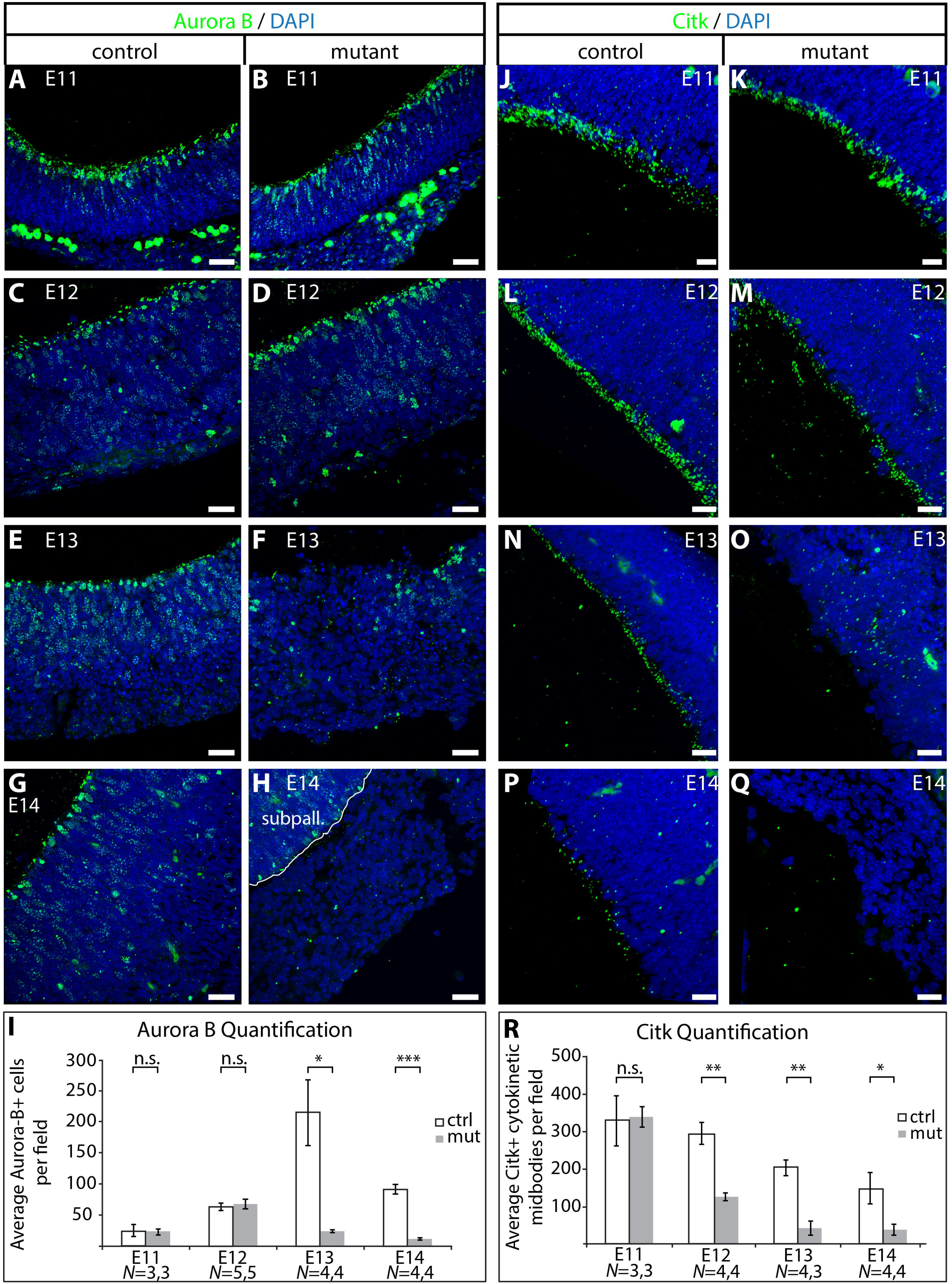
U11 loss results in cell cycle defects. (**A-H**) IF for Aurora B (green) on sagittal sections of the control (left panels) and mutant (right panels) pallium across E11, E12, E13, and E14. Nuclei are marked by DAPI (blue). (**I**) Bar graph showing quantification of the average number of Aurora-B+ cells in a 240 μm^2^ field in the control and mutant pallium. (**J-Q**) IF for Citk (green) on sagittal sections of control (left panels) and mutant (right panels) pallium across E11, E12, E13, and E14. (**R**) Bar graph showing quantification of the average number of Citk+ cytokinetic midbodies per 240 μm^2^ field in the control and mutant pallium. For quantifications, statistical significance was determined by two-tailed student’s t-tests. N.s.=not significant; *=*P*<0.05; **=*P*<0.01; ***=*P*<0.001. Scale bars=30 μm.

## Discussion

Here, we report the role of minor splicing in cortical development by *Emx1*-Cre-mediated U11 snRNA ablation. While the mutation underlying MOPD1 and Roifman syndrome is in the U4atac snRNA and our mouse model targets U11, both lead to inactivation of the minor spliceosome (Fig. 4D1) (Edery et al., 2011; He et al., 2011; Merico et al., 2015). Thus, the microcephaly in *Rnu11* mutant mice corroborates the connection between minor spliceosome inactivation and microcephaly in MOPD1 and Roifman syndrome (Edery et al., 2011; He et al., 2011; Merico et al., 2015). Moreover, our findings shed light onto the potential molecular and cellular defects contributing to the microcephaly observed in MOPD1 and Roifman syndrome.

Cell death is the primary cause of the microcephaly in the *Rnu11* cKO mice (Fig. 2K-Y, Fig. S6A-G). This death was first observed at E12 (Fig. 2R-S, Fig. S6A-B, S6G) whereas U11 loss began at E10 (Fig. 2B), suggesting that U11-null cells can survive for two days without minor splicing. The survival of U11-null cells might be due to the presence of previously spliced MIG transcripts, and their loss is finally experienced at E12, due to turnover (Wilusz et al., 2001). Another possibility is that U11 is required in a cell-type specific manner. Indeed, we found that loss of U11 predominantly affected the RGC population (Fig. 3N, Fig. S4C). In contrast, there was no significant change in the number of IPCs between E12 and E13 mutant pallium (Fig. S4D). The majority of IPCs undergo symmetric neurogenic divisions, to produce two neurons (Noctor et al., 2004). In the absence of replenishment of the IPC population, symmetric neurogenic divisions would deplete this population. The lack of IPC depletion at E13 (Fig. S4D) suggests that the IPC population was being actively maintained in the E13 mutant pallium. IPCs are produced from either asymmetrically dividing RGCs or IPCs that undergo symmetric proliferative divisions (Huttner and Kosodo, 2005). Therefore, maintenance of the IPC population in the E13 mutant pallium can occur through one of two paths: (1) U11-null RGCs divide asymmetrically to produce IPCs, or (2) a higher percentage of U11-null IPCs undergo proliferative divisions than observed in normal development. The latter case would be expected to reduce the number of neurons in the E13 pallium, relative to the control. However, we found no significant change in the number of neurons in the E13 mutant and control pallium (Fig. S4A), thereby supporting the former possibility. From our data, it is unclear whether the U11-null RGCs undergo asymmetric division to produce neurons, whether the IPCs themselves undergo apoptosis, or whether IPC loss is due to depletion by symmetric neurogenic division (Fig. 6). In summary, inactivation of the minor spliceosome in the U11-null developing mouse cortex first results in the death of self-amplifying RGCs (Fig. 6). However, RGCs undergoing asymmetric divisions to produce IPCs survive (Fig. 6). Since each asymmetrically dividing RGC only produces one RGC daughter cell, these divisions cannot offset the loss of self-amplifying RGCs (Fig. 6). Moreover, in the mutant pallium, fewer IPCs are produced as the number of RGCs declines, eventually resulting in IPC depletion (Fig. 6). Ultimately, the mutant pallium is comprised primarily of neurons by E14 (Fig. 6).

**Figure 6.**
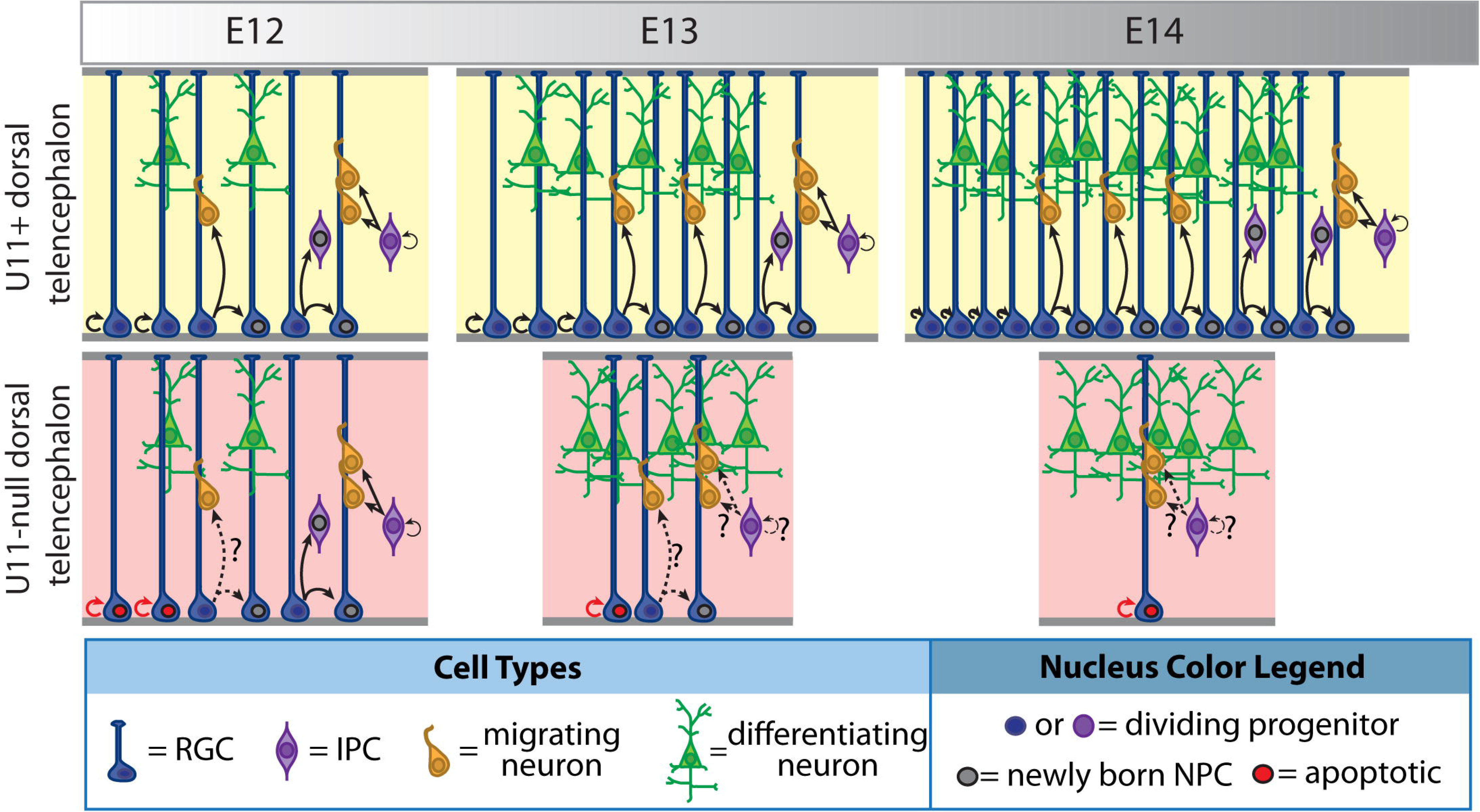
Model of cortical development in *Rnu11* mutant mice. Schematic representing cortical development in normal (top, yellow background) and mutant (bottom, pink background) mice, from E12 to E14. Radial glial cells (RGCs) are shown in blue, intermediate progenitor cells (IPCs) in purple, migrating neurons in orange, and differentiating neurons in green. Black arrows show cell division events. Cells undergoing cell death are indicated by red nuclei/arrows. Gray nuclei mark newly born neural progenitor cells (NPCs) produced from divisions at the specific time-point shown. Question marks and dashed lines indicate events that remain to be explored.

The requirement of minor splicing for rapidly dividing cells is reflected by the early embryonic lethality observed after constitutive *Rnu11* ablation (Fig. 1G). Similar observations of embryonic lethality have been reported in *Drosophila* and zebrafish, further underscoring the importance of proper MIG expression in cycling stem cells (Otake et al., 2002; Markmiller et al., 2014). Even within a progenitor pool in the developing cortex, we observed vastly different outcomes of U11 loss, which are dependent on both progenitor cell type (RGC vs IPC) and the type of cell division (Fig. 6). For example, U11-null RGCs were able to produce IPCs, while U11-null RGCs undergoing self-amplifying divisions were the first to be depleted in the U11-null pallium (Fig. 3N, Fig. 6, Fig. S4C&D). Notably, our data suggest that U11-null neurons are also resistant to U11 loss, since (1) there was no significant change in neuron number in the mutant pallium between E13 and E14 (Fig. S4B), and (2) >95% of the cells remaining in the E14 U11-null pallium were neurons (Fig. 3G). Together, these findings indicate a cell type-specific susceptibility to U11 loss, with rapidly cycling, self-amplifying RGCs being the most vulnerable and post-mitotic neurons the least (Hardwick et al., 2015). This cell type-specificity might also explain why constitutive, 90% loss of minor spliceosome activity in MOPD1 patients predominantly affects development of the cortex and limbs, while the development of other tissues is unaffected (He et al., 2011).

RNAseq analysis revealed that overall minor intron retention was elevated in the E12 mutant pallium compared to the control (Fig. 4D1). While this finding is consistent with the RNAseq performed on samples from patients with disrupted minor intron splicing, we found that the change in the median value of minor intron retention between the control and mutant samples was lower than reported by these groups (Fig. 4D1) (Madan et al., 2015; Merico et al., 2015). It is possible that this difference reflects tissue specificity. Another variable in our data is the presence of U11+ cells in the tissue dissected for RNAseq analysis (Figs 2F-F’, 4A inset). Finally, unlike patient samples that have systemic, long-term loss of minor splicing activity, we harvested the tissue for RNAseq analysis two days post-Cre-mediated U11 ablation (Fig. 2B&F). Therefore, the persistence of previously spliced MIG-transcripts due to turnover kinetics in our system could contribute to the shift in the median value (Wilusz et al., 2001). Similar experimental issues could account for the variations in minor intron retention that have been reported in HeLa cells treated with U6atac morpholino (Younis et al., 2013). Nevertheless, 40% of minor introns showed significantly elevated minor intron retention in the mutant (Supplementary Data File 2). In most of these cases, one would expect that minor intron retention would activate the nonsense-mediated decay pathway, resulting in downregulation of MIGs. However, 168 out of the 169 MIGs with significantly elevated minor intron retention in the mutant did not show the corresponding significant downregulation at the level of gene expression (Supplementary Data File 1&2). This is in agreement with the finding that mutation in an essential minor spliceosome component (*rnpc3*) in zebrafish did not affect expression levels of 90% of the MIGs, while still resulting in elevated minor intron retention (Markmiller et al., 2014). These findings suggest that inactivation of the minor spliceosome, while disrupting minor intron splicing, does not necessarily result in downstream changes in overall gene expression.

There were 20 MIGs involved in cell cycle regulation that showed increased minor intron retention, including the kinetochore assembly gene *Spc24*, which was also downregulated in the mutant (Fig. 4C&G, Table S2) (McCleland et al., 2004). Moreover, our observation that reduction in the RGCs undergoing cytokinesis precedes NPC loss indicates that minor splicing loss results in cell death due to cell cycle defects (Figs 3G, 5R). This finding, in combination with the observations that rapidly amplifying stem cells in early embryonic development and rapidly dividing, self-amplifying RGCs are predominantly affected by U11 loss (Figs 1G, 3N, S4C), suggests a connection between cell cycle speed and sensitivity to inactivation of the minor spliceosome (Arai et al., 2011; Borrell and Calegari, 2014). In addition to the disruption in cell cycle-regulating MIGs, our RNAseq analysis revealed significant upregulation of p53-activated, pro-apoptotic genes (Table S1). Given that none of these genes contain minor introns, their upregulation is likely a secondary effect, caused by minor intron retention in MIGs functioning upstream of p53 activation, such as in DNA damage response (Williams and Schumacher, 2016). This is supported by the finding that 13 MIGs with elevated minor intron retention enriched for the DAVID function “response to DNA damage stimulus” (Fig. 4E). Moreover, manual curation of the functions performed by the 169 MIGs with significantly elevated minor intron retention revealed an additional 4 MIGs involved in various steps of the DNA damage response pathway (Table S3). Together, our RNAseq findings indicate that the observed RGC death in the U11-null pallium is caused by both cell cycle defects and p53 activation, likely due to DNA damage accumulation. However, given that we identified 169 MIGs with significantly elevated minor intron retention, and that these MIGs have varied functions, it would be shortsighted to conclude that these defects are the only causes of the NPC death in the mutant pallium. It is likely that disruption of multiple MIGs, and therefore disruption of their varied functions, contribute to the observed NPC death. To identify a short-list of MIGs whose dysfunction may underlie this death, we interrogated the 169 MIGs in recently published essentialome reports, which identified genes essential for cycling cell survival in various cancer cell lines (Blomen et al., 2015; Hart et al., 2015; Wang et al., 2015). Of the 169 MIGs with increased minor intron retention, 17 MIGs are common to all the cycling cell essentialomes (Table S4). Even within this short-list of genes, these MIGs execute various functions, including cell cycle (6 genes), RNA processing (7 genes), transcription regulation (3 genes), and translation (3 genes) (Table S4). Future investigation will both allow us to understand the impact of loss of function of these MIGs on the observed NPC loss in the U11-null pallium, and provide insight into why neurons and NPCs undergoing differentiative divisions are less susceptible to U11 loss.

## Materials and Methods

### Animal Procedures and Generation of the *Rnu11* cKO Mouse

All mouse procedures were performed according to the protocols approved by the University of Connecticut Institutional Animal Care and Use Committee. The *Rnu11* conditional knockout mouse was generated by the University of Connecticut Health Center, using a vector designed to target the *Rnu11* locus by homologous recombination. This construct contained a loxP site 1090 bp upstream of *Rnu11*, an Frt-flanked phosphoglycerine kinase (PGK)-Neo cassette, a loxP site 1159 downstream of *Rnu11*, and a PGK-dTA (diphtheria toxin A) negative selection cassette downstream of the 3’ arm of homology (Fig. S1). This construct was targeted into 129X1/SvJ mouse embryonic stem (ES) cells, followed by validation of successful recombination (SI Methods). *EIIa*-Cre and *Emx1*-Cre were bred into the *Rnu11* conditional knockout line to target *Rnu11* for germline ablation and for removal in the developing forebrain, respectively (Lakso et al., 1996; Gorski et al., 2002). All experiments were performed using *Rnu11^WT/Flx^* and *Rnu11^Flx/Flx^* mice that were *Emx1*-Cre^+/-^.

### PCR

Either 25 ng of gDNA or cDNA prepared from total RNA extracted from control (*N*=3) and mutant (*N*=4) E12 pallium were used for PCR, as described in the SI Methods.

### Transcardial Perfusion

P70 control and mutant mice were anesthetized using isoflurane, followed by transcardial perfusion performed as previously described (Gerfen, 2003). After the brains were removed, they were Flxed overnight in 4% PFA at 4°C, followed by PBS washes and cryoprotection in 30% sucrose in PBS at 4°C for two days. Following an overnight incubation in half 30% sucroselhalf OCT compound (Fisher Scientific, #23-730-571), the brains were embedded in OCT compound and sectioned using a cryostat (Leica CM 3050 S).

### *In Situ* Hybridization

16 μm cryosections of mouse heads (for E10–E12 and P0), telencephalons (E13–E14), and whole brains (for P21), *N*=3 for each time-point and genotype, were used for section *in situ* hybridization (SISH). SISH was performed using antisense, digoxigenin-labeled U11 RNA probe, which was detected using either alkaline phosphatase (AP) or fluorescent labeling (FISH), as described (Baumgartner et al., 2014). The U11 probe was generated using the U11 expression primers in Table S5.

### TUNEL Assay

16 μm cryosections of either embryonic mouse heads (for E10 – E12) or telencephalons (E13 – E14) were used for terminal deoxynucleotidyl transferase dUTP nick ending labeling (TUNEL) analysis. The *in situ* cell death detection kit, TMR red (Roche, #12156792910), was used, according to the manufacturer’s instructions.

### Immunohistochemistry

16 μm cryosections of either mouse heads (for E10 – E12), telencephalons (E13 – E14), or whole brains (P0 and P70) were used for hematoxylin and eosin staining and immunofluorescence experiments, as described in the SI Appendix (SI Methods).

### Image Acquisition and Quantification

Processed slides were imaged with a Leica SP2 confocal microscope, where settings for laser intensity, excitation-emission windows, gain, and offset conditions were identical between control and mutant sections on a given slide. For every slide that was processed for ISH, FISH, or IF, the slide contained at least one control and one mutant section. Sections serial to those that revealed loss of U11 by ISH analysis were used for IF staining, and images were collected in regions of the pallium that showed U11 loss in the mutant by ISH (Fig. 2D, 2F, 2H, 2J). For each channel for every experimental condition, confocal imaging settings were optimized for fluorescence in the control section on a given slide, and maintained for all other sections on that slide. For TUNEL imaging, collection settings were adjusted using the control section. These settings were maintained when imaging the mutant section(s) on the same slide. Further processing was performed on IMARIS v8.3.1 (Bitplane Inc.) and Adobe Photoshop CS4 (Adobe Systems Inc.). Overall, all image processing in IMARIS and Photoshop was identical between images of control and mutant sections from the same slide. Manual quantification was performed using the spot tool in the IMARIS software; statistical significance was calculated using two-tailed student’s *t*-tests. For analysis of NeuN+, Pax6+, and Tbr2+ cell quantification across mutant cortical development, statistical significance was determined by one-way ANOVA, followed by the post hoc Tukey test.

### Bioinformatics Analysis

Total RNA was extracted from control (*N*=5) and mutant (*N*=5) E12 pallium. Total RNA libraries were prepared by Illumina TruSeq Stranded Total RNA Library Sample Prep Kit (RS-122-2201) with Ribozero to remove ribosomal RNA, and were then sequenced using Illumina NextSeq 500, which generated 151 bp reads. Reads were mapped against the mm10 genome using Hisat2 (Kim et al., 2015). Gene expression was determined using ISOEM2 and ISODE2 (Nicolae et al., 2011; Al Seesi et al., 2014). BEDTools was used for intron retention analysis (Fig. S7, SI Methods) (Quinlan and Hall, 2010).

### RNA Extraction and cDNA Preparation

Palliums were dissected from E12 *Rnu11* control (*N*=3) and mutant (*N*=4) mice, and the tissue collected from each embryo was individually used for total RNA isolation. Tissue from each embryo was separately triturated in 100 μL of TRIzol (Invitrogen, #15596026). RNA was extracted using the DirectZOL^™^ RNA MiniPrep kit (Zymo Research, #R2050), per the manufacturer’s instructions. For total RNA samples, 500 ng of RNA was used for cDNA synthesis, which was performed as previously described (Kanadia et al., 2006).

## Acknowledgements

We would like to thank Drs. Anastasios Tzingounis and Heun Soh for sharing the *Emx1*-Cre line, Dr. Akiko Nishiyama for sharing the *EIIa*-Cre line, and Dr. Alexander Jackson for sharing the Ai9 tdTomato Cre reporter line. Moreover, we would like to thank Dr. Siu-Pok Yee from the Gene Targeting and Transgenic Facility at the UConn Health Center for generating the *Rnu11* conditional knockout mouse, and Dr. Bo Reese from the Institute for Systems Genomics Center for Genome Innovation for performing RNA sequencing. Finally, we would like to acknowledge Dr. Ion Mandoiu for his assistance with bioinformatics analyses, and Mr. Ethan Cope for his help with microscopy.

